# Expanding discovery from cancer genomes by integrating protein network analyses with in vivo tumorigenesis assays

**DOI:** 10.1101/151977

**Authors:** Heiko Horn, Michael S. Lawrence, Candace R. Chouinard, Yashaswi Shrestha, Jessica Xin Hu, Elizabeth Worstell, Emily Shea, Nina Ilic, Eejung Kim, Atanas Kamburov, Alireza Kashani, William C. Hahn, Joshua D. Campbell, Jesse S. Boehm, Gad Getz, Kasper Lage

## Abstract

Approaches that integrate molecular network information and tumor genome data could complement gene-based statistical tests to identify likely new cancer genes, but are challenging to validate at scale and their predictive value remains unclear. We developed a robust statistic (NetSig) that integrates protein interaction networks and data from 4,742 tumor exomes and used it to accurately classify known driver genes in 60% of tested tumor types and to predict 62 new candidates. We designed a quantitative experimental framework to compare the *in vivo* tumorigenic potential of NetSig candidates, known oncogenes and random genes in mice showing that NetSig candidates induce tumors at rates comparable to known oncogenes and 10-fold higher than random genes. By reanalyzing nine tumor-inducing NetSig candidates in 242 patients with oncogene-negative lung adenocarcinomas, we find that two (*AKT2* and *TFDP2*) are significantly amplified. Overall, we illustrate a scalable integrated computational and experimental workflow to expand discovery from cancer genomes.

## Introduction

Cancers are initiated when somatic mutations, copy number alterations, or genomic fusion events of specific genes (hereafter, we will refer to these as driver genes or cancer genes) confer a selective advantage to the corresponding cell thus promoting tumorigenesis. Identifying the driver genes in tumors of individual cancer patients provides key diagnostic and therapeutic insight. Therefore, it is a central aim of oncology to provide a complete catalogue of genes underlying human cancers^1–6^.

The recent revolution of cancer genome analyses has enabled the statistical identification of cancer genes in an unbiased manner based on somatic mutations or copy number changes using gene-based statistical tests such as MutSig, Oncodrive, GISTIC and RAE^7–10^. These methods have identified many genes mutated or amplified at high frequencies (>20%) in tens of tumor types^11^. However, for many tumor types, insufficient sample numbers, compounded by high background mutation and copy number rates^11^ render it challenging to confidently pinpoint driver genes at intermediate (2-20%) or low (<2%) frequencies. For this reason, a large number of putative driver genes don't meet established statistical cutoffs, and a large number of biologically or clinically relevant driver genes remain to be discovered.

Many existing methods have been used to highlight network modules (based on functional genomics data) that are significantly mutated in tumors^12–18^. These analyses have been valuable for illuminating the biological processes and pathways involved in cancers (reviewed in Creixall et al.^19^). However, the evidence from network-based approaches comes from aggregating weak genetic signals in a set of genes connected into a network and not from an overwhelming mutation signal in any individual gene itself. This means that there is no strong direct link between specific genes in a significantly mutated module and the cancer in question. Additionally, since most network-based methods are only retrospectively evaluated through benchmarks, or by rigorously following up on only a few new genes, it is impossible determine how much ‘knowledge contamination’ (i.e., the notion that genes are more studied because they are cancer genes) introduces circularity and biases the outcomes towards more well established, or classic, cancer networks. These issues could be addressed by executing a systematic large-scale comparison of the tumorigenic potential of tens of genes embedded in a significantly mutated network versus a large number of known cancer genes (positive controls) and random genes (random controls), but such an analysis is lacking. Together this means that, if the aim is specifically to expand cancer gene discovery amongst genes with inconclusive mutations in existing cancer genomes, the real predictive value of network-based approaches remains unclear.

Towards this aim, we developed a statistic (NetSig) that combines cancer mutation data and molecular network information to expand cancer gene discovery from tumor genomes. With NetSig we had a particular focus on addressing the effects of ‘knowledge contamination’ and on designing a method that is independent of gene-based statistical tests like MutSig, Oncodrive, GISTIC and RAE so that it can seamlessly complement these approaches in any tumor genome analysis pipeline. To test the predictive power of NetSig, we developed an *in vivo* quantitative experimental framework that enabled us to compare the tumorigenic potential of 23 genes with a significant NetSig score to that of 25 known cancer genes and 79 random genes in mouse experiments. Based on the network analysis and *in vivo* experiments nine candidates were particularly relevant to lung adenocarcinoma. We reanalyzed copy number data derived from 660 patients with lung adenocarcinoma to discover higher rates of amplification of *TFDP2* and *AKT2* in patients without established genomic driver events compared to patients with mutations and amplifications in known oncogenes. The code is available from www.lagelab.org/resources and the algorithm has been implemented din FireCloud (https://software.broadinstitute.org/firecloud/).

## Results

### Design and properties of the NetSig statistic

NetSig, combines data from existing cancer genomes (spanning 21 tumor types and 4,742 tumor genomes) and InWeb (a human protein-protein interaction network that has been used in the 1000 Genomes Project^20^) to calculate the mutation signal in a genes’ functional protein-protein interaction network. Since we specifically wanted to test the predictive power of mutations in a gene’s network, we excluded mutation information on the gene itself in the calculation of the NetSig statistic (for all details see **Methods**).

To benchmark NetSig and to understand the effect of ‘knowledge contamination’ on the statistic, we defined a set of ‘Cosmic classic’ (i.e., very well established) cancer genes from the Cosmic database (http://cancer.sanger.ac.uk/cosmic) and a set of ‘recently emerging cancer genes’ from recent sequencing studies (**Methods**, **Supplementary Table 1**). To test for cryptic confounders we also defined a set of random genes (**Methods**, **Supplementary Table 1**). We confirmed that the ‘Cosmic classic’ and ‘recently emerging’ sets can be classified based on their NetSig score with an area under the receiver operating characteristics curve (AUC) of 0.86, and 0.75, respectively (**Fig. 1a**, adj. *P* < 0.05 for each of these AUCs, using permuted networks, **Supplementary Figure 1**). As expected the random control genes fit the null hypothesis and cannot be distinguished from other genes represented in InWeb (**Fig. 1a**, AUC 0.49, *P* = 0.8). We further show that NetSig can accurately classify cancer genes in ~ 60% of the tumor types for which we have data (**Methods** and **Supplementary Note 1**, **Supplementary Figure 2**), illustrating the potential of our statistic to inform many different individual tumor types.

**Figure 1.**
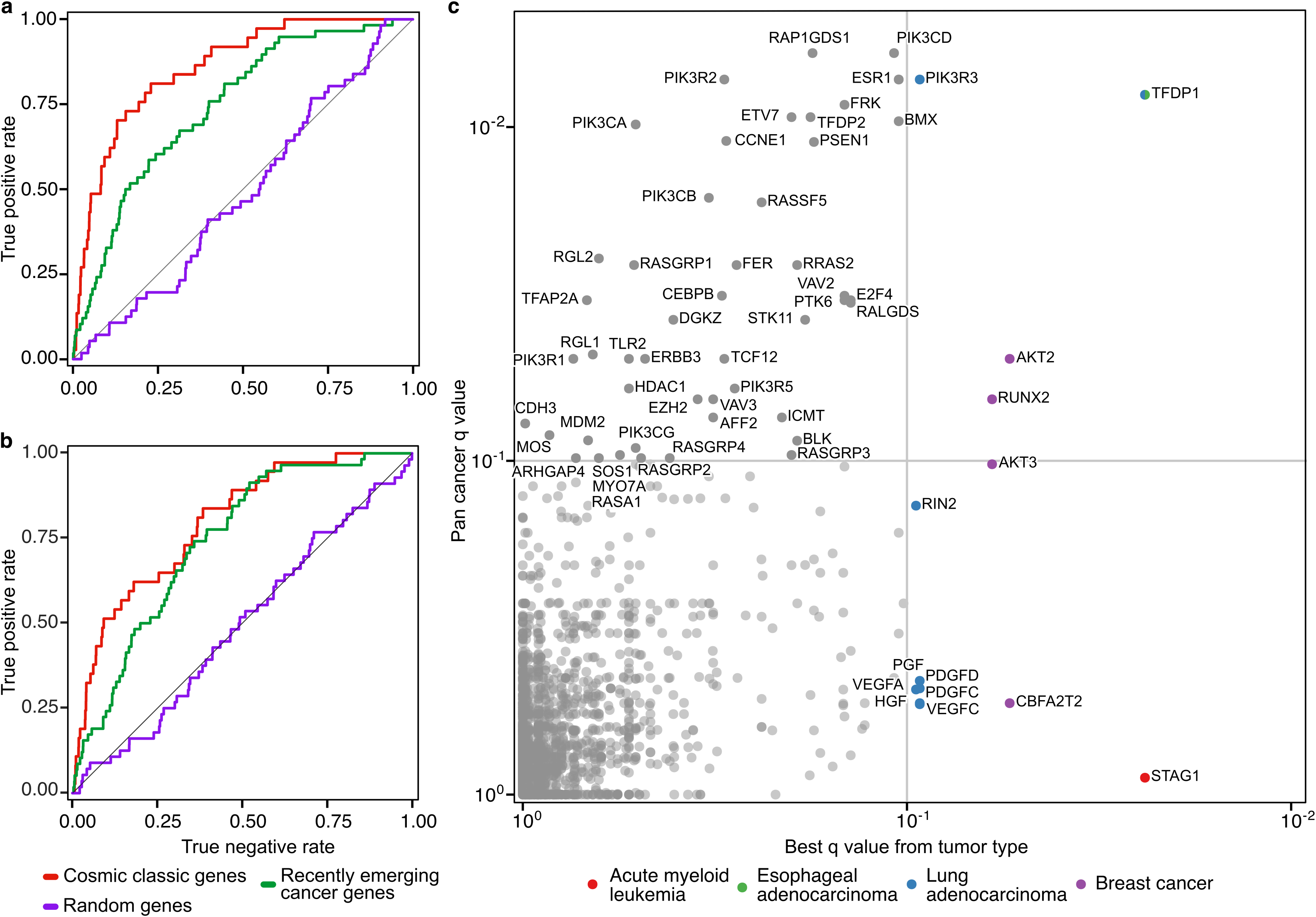
NetSig predicts true cancer genes. **a)** Areas under the receiver operating characteristics curve (AUCs) for genes in the ‘Cosmic classic’ and ‘recently emerging’ sets are 0.86 and 0.75, respectively (adj. *P* < 0.05). Genes from the random set fit the null hypothesis (AUC 0.49, nominal *P* = 0.75). **b)** AUCs when removing the effect of very well established cancer genes is 0.79, 0.73, and 0.5, for the “Cosmic classic”, “recently emerging”, and random sets, respectively. **c)** Visualizing the NetSig500 set. Genes are represented as individual dots and plotted along the x-axis by the NetSig Q value from the most significant of 21 tumor types, and on the y-axis by the NetSig Q value when 4,724 tumors are analyzed as a combined pan-cancer cohort. Significance at FDR Q <= 0.1 is indicated on each axis by grey lines.

The majority of genes scored by NetSig fit the null hypothesis and lie on the diagonal in a quantile-quantile plot, but there is an overall genomic inflation (lambda = 1.29) of the significances assigned to genes (**Supplementary Figure 3**). This could be due to ‘knowledge contamination’, the inherent polygenic nature of cancers, or a combination. To dissect this phenomenon, we removed the effect of well-studied cancer genes from our analysis (**Methods**, **Supplementary Note 2**). The ability to predict ‘Cosmic classic’ cancer genes is reduced (**Fig. 1b**, from an AUC of 0.86 to 0.79) indicating some ‘knowledge contamination’ of this set, but effect on ‘recently emerging’ cancer genes is much less pronounced (**Fig. 1b**, from an AUC of 0.75 to 0.73). Consistent with these observations, we see that removing the effect of the ‘Cosmic classic’ gene set reduces the lambda from 1.29 to 1.09 in the quantile-quantile plot and that it only changes slightly to 1.07 when the impact of the ‘recently emerging’ set is also removed (**Supplementary Figure 3**). Furthermore, running NetSig on random networks results in a non-inflated quantile-quantile plot as expected (**Supplementary Figure 3**). We also show that NetSig adequately normalizes for the amount of interactions a gene has at the protein level (**Supplementary Figure 4**).

Together, these analyses show that there is some ‘knowledge contamination’ in the protein-protein interaction data specifically of the ‘Cosmic classic’ set and that this leads to a significantly inflated AUC in the benchmark if it is not taken into consideration. Conversely, there is almost no ‘knowledge contamination’ of genes emerging from recent sequencing studies. This means that when predicting new cancer genes from existing cancer genomes, ‘knowledge contamination’ should not confound the NetSig method when applied to protein-protein interaction data from InWeb.

### Predicting NetSig candidates from 4,742 tumor genomes

To test if NetSig can predict new likely driver genes from existing cancer genome data we calculated NetSig scores of all genes that had at least one high-confidence protein interaction in InWeb and adjusted the result of this analysis for multiple hypothesis testing. We calculated NetSig scores both using the pan-cancer cohort of 4,742 tumors and using mutation data from each of the individual 21 tumor types represented in^11^ (**Methods**). We declared genes with a false discovery rate (FDR) Q ≤ 0.1 using the pan-cancer data significant (**Fig. 1c**) and also declared genes with a Q ≤ 0.1 in each of the individual 21 tumor types significant.

The pooled set (named NetSig5000, **Supplementary Table 2**) contains all unique genes that were significant in the pan-cancer analysis or in at least one of the 21 tumor types. Our NetSig5000 set comprises 62 genes, of which we divided into five groups based on their known connection to cancer. Groups 1 (n = 12) and 2 (n = 9) contain genes already known to be involved in cancers based on significant point mutations or gene fusion events, respectively. These groups serve as a positive control that NetSig can identify known cancer genes. Groups 3 (n = 24) and 4 (n = 13) contain genes that have been speculated to be causal in cancers based on evidence from model systems or from gene expression analyses. Group 5 (n = 4) have never been linked to cancer (see **Supplementary Table 3** and **Supplementary Note 3** for more information about genes in the NetSig5000 set and **Supplementary Figure 5** for examples of NetSig networks). All results can be accessed and visualized at http://www.lagelab.org/resources/.

### Comparing the tumorigenic potential of 23 NetSig candidates, 25 oncogenes, and 79 random controls

To confirm whether NetSig nominates candidate genes with bona fide tumorigenic potential, we tested the tumorigenic potential of 23 genes from the NetSig5000 set (**Supplementary Table 4**, for selection criteria of these 23 genes see **Methods**), 79 different patient-derived mutations of 25 known driver genes (positive control, **Supplementary Table 5**) and 79 random genes (random control, **Supplementary Table 6**) using a massively parallel *in vivo* tumorigenesis assays (**Fig. 2a** and **Methods**). The assay transduces and overexpresses barcoded cDNA constructs of candidate genes (and alleles representing patient-derived mutations) into activated small-airway epithelial cells [SALE-Y cells^21^] or activated immortalized kidney epithelial cells [HA1E-M cells^21–24^]. For each cell model (SALE-Y or HA1E-M) all genes (or alleles) to be tested are pooled, grown, and injected subcutaneously into immunocompromised animals at three injection sites per animal. In animals that develop tumors, driver genes can be identified by homogenizing tumors in the animals and by sequencing the barcodes found in the tumor cells (**Methods**).

**Figure 2.**
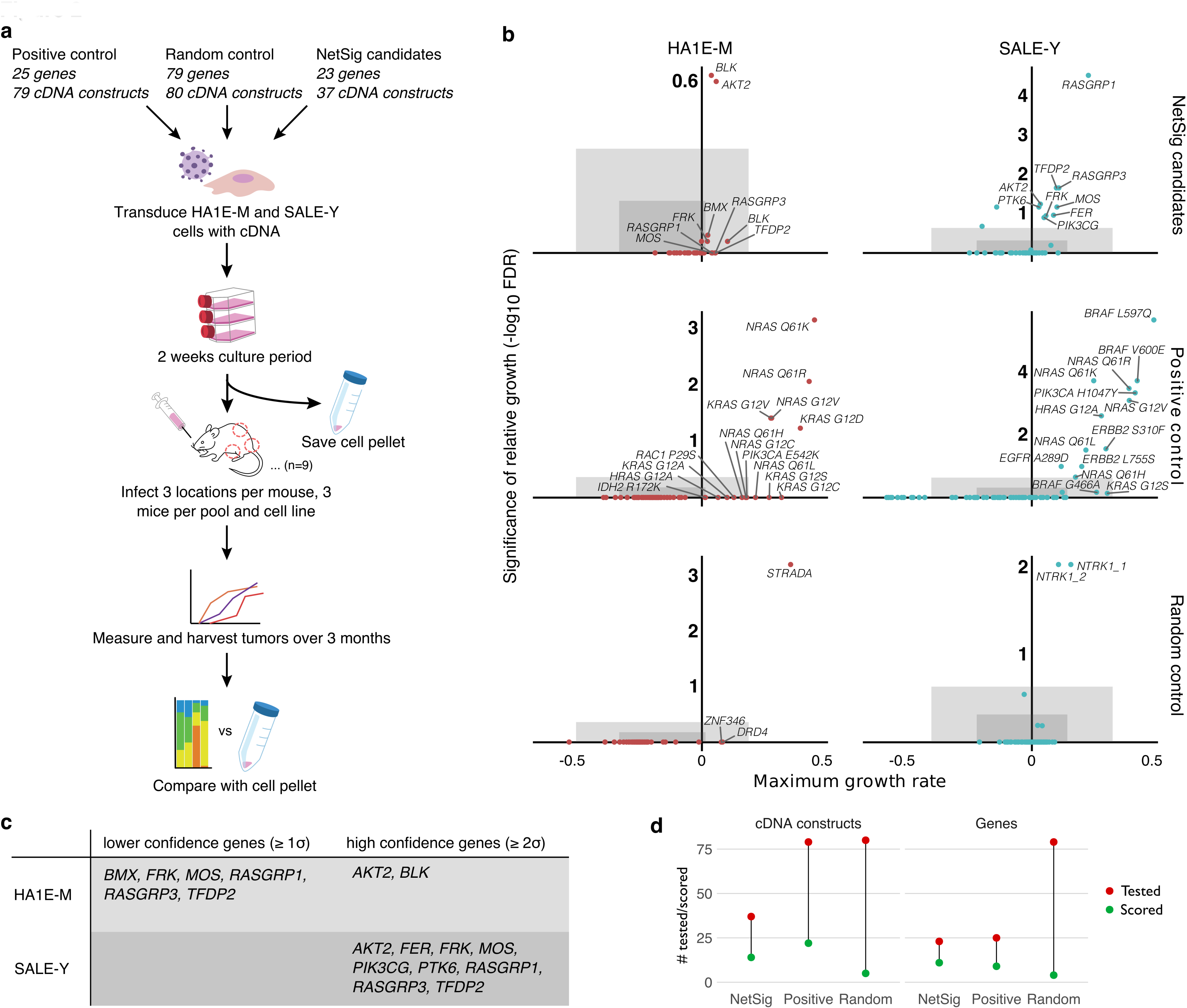
*In vivo* tumor formation of NetSig5000 and control sets. **a)** Experimental design. **b)** Tumorigenic potential of 23 NetSig5000 genes (NetSig candidates), 25 known oncogenes (Positive control), and 79 random genes (Random controls) in *in vivo* mouse tumorigenesis experiments. X-axis indicates maximum proliferation rate and y-axis maximum significance of enrichment in tumors relative to pre-injection samples. Dark grey boxes indicate one standard deviation from the median (lower confidence) and light grey boxes two standard deviations from the mean (higher confidence). **c)** Candidates that induce tumors at the higher and lower confidence threshold stratified by cell model. **d)** Proportion of the NetSig5000 candidates, positive control set, and random set, respectively, that induced tumors in mice. Left panel indicate the results at the level of cDNA constructs. Right panel indicate results at the gene level.

To compare the tumorigenic potential of the three gene sets across multiple cell models we developed a quantitative analytical framework that defines a gene as tumorigenic based on both *in vivo* proliferation rate of the tumor cells and the significance of the relative growth of the tumors (**Methods**). Our analysis showed that many of the tested NetSig5000 genes (11/23 or 48%) are indeed capable of driving tumorigenesis (**Fig. 2b** and **c, Supplementary Table 7**). Specifically, these pooled screening results support the putative tumorigenic potential of *AKT2, BLK*, *BMX*, *FER*, *FRK*, *MOS*, *PIK3CG, PTK6, RASGRP1, RASGRP3*, and *TFDP2*),for a comprehensive literature review of these genes see **Supplementary Notes 4**). In comparison, the proportion of known driver genes from the positive control set that induced tumors was 9/25 or 36%, providing an estimate of assay sensitivity, (**Fig. 2d**, see **Methods** for details) and the proportion of random genes that induced tumors was 4/79 or 5%, providing an estimate of the assays false positive rate (**Fig. 2d**). We note that the two random genes that induced tumors are *NTRK1* (encoding a tyrosine kinase with established tumorigenic properties) and *STRADA* (an interactor of the *STK11* tumor suppressor at the protein level), suggesting that these could be real driver genes that remain to be discovered, that the assay has false positive rate well below 5%, meaning that the specificity of the assay is likely to be above 95%.

### Significant copy number gains of *TFDP2* and *AKT2* in lung adenocarcinoma patients

Nine NetSig5000 genes (*AKT2, FER*, *FRK*, *MOS*, *PIK3CG, PTK6, RASGRP1, RASGRP3*, and *TFDP2*) validated with high confidence in a cell model (SALE-Y) that is particularly relevant for exploring genes that can induce lung adenocarcinomas^21^. Based on this observation we hypothesized that a subset of these nine genes may be responsible for driving lung adenocarcinomas in oncogene negative patients [meaning patients that do not have a known oncogenic driver event in the RAS/RAF/receptor tyrosine kinase (RTK) pathway as previously described^25^].

To test these hypotheses we used a dataset of 660 lung adenocarcinomas from TCGA and related studies^25–27^. We first tested for copy number differences between the oncogene negative (n = 242) and the oncogene positive patients (n = 418), showing that the nine genes as a set have a significantly higher copy number in the former group (*P* = 7.0e-3, using Fisher’s exact test, **Fig. 3a**). We also show that *TFDP2* and *AKT2* are individually found at higher copy numbers in the oncogene negative group (FDR < 0.1 for each gene, **Fig. 3a**).

**Figure 3.**
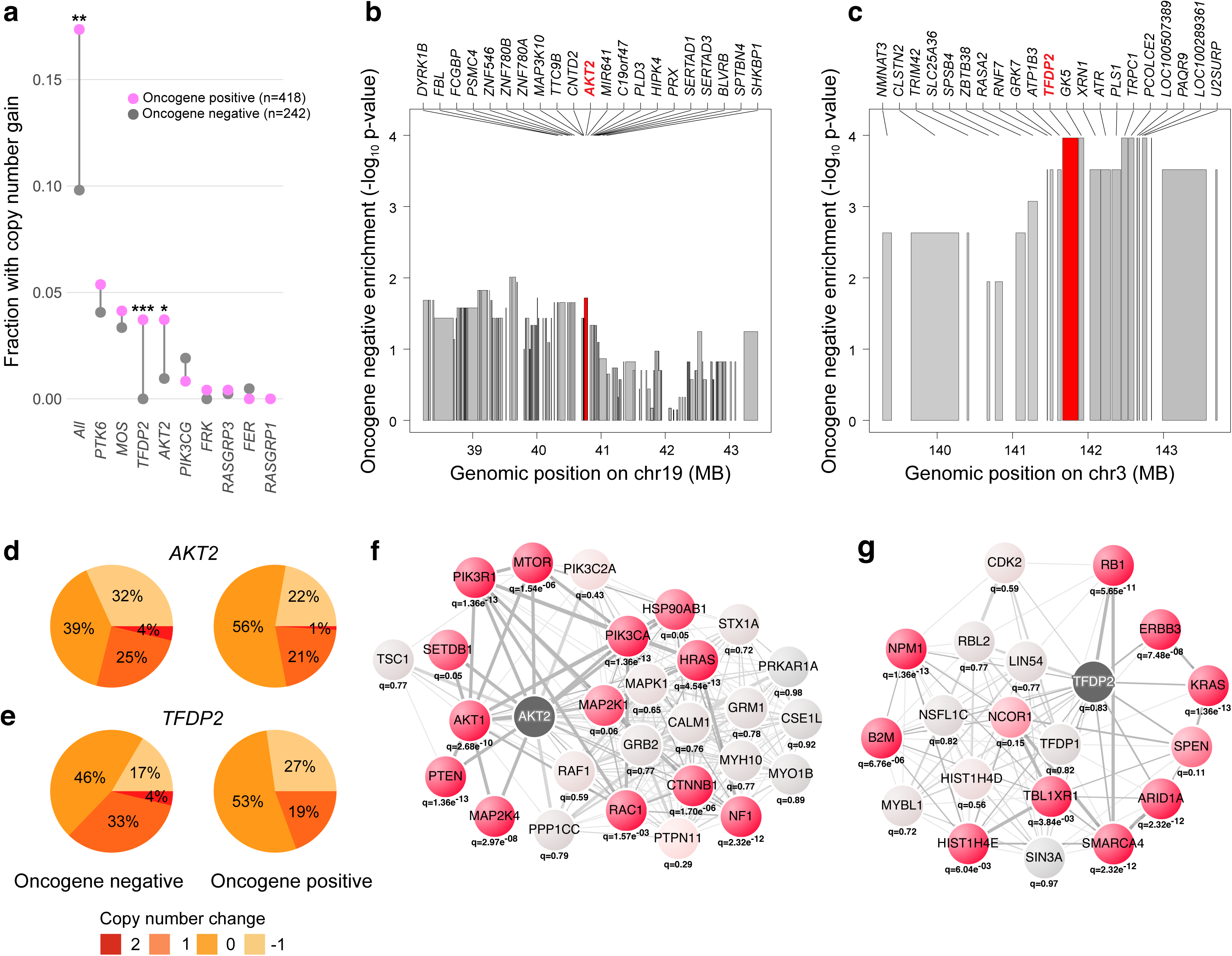
Targeted re-analysis of oncogene negative lung adenocarcinoma patients. **a)** Amplification of the nine genes that induce tumors in the lung adenocarcinoma-relevant cell model. As a group the genes are significantly amplified (*P* = 7.0e-3) and *AKT2* and *TFDP2* are individually significantly amplified (FDR Q < 0.1). **b**), **c)** In depth view of the amplified regions surrounding *AKT2* and *TFDP2*, respectively. **d)**, **e)** The proportion of oncogene positive or negative patients with −1, 0, 1, or 2 copy number changes of *AKT2* or *TFDP2*. **f)**, **g)** NetSig networks of *AKT2* and *TFDP2*. Nodes other than *AKT2* and *TFDP2* are colored by the significance of the pan-cancer Q value of the corresponding gene, where light grey represents Q close to 1 and red Q << 1, with darker red representing more significant Q values as indicated below the relevant node.

Through an in-depth analysis of the surrounding genomic regions, we ruled out that adjacent potential oncogenes are underlying the signal we see (**Figs 3b** and **c** and **Supplementary Note 5**) and we also confirmed that there is no overall difference in copy numbers between the two patient groups (**Supplementary Figure 6**). The genomic events observed for *AKT2* and *TFDP2* are not high-level amplifications. Rather, 3 and 4% percent of the oncogene negative patients have two extra copies of *AKT2* and *TFDP2*, respectively; and 4 and 14% have one extra copy of *AKT2* and *TFDP2*, respectively (**Figs 3d** and **e**). In the nine genes we also tested whether there is evidence for increased rates of gain-of-function single nucleotide variants (SSNVs) or insertions or deletions (indels) in the oncogene negative versus positive group, which is not the case.

Given the dominating effect of the RAS/RAF/RTK pathway in lung adenocarcinoma, a more straightforward approach to gene discovery would be to make a targeted analysis of mutations or copy number gains in genes in the extended RAS/RAF/RTK pathway (defined here as genes that have at least one protein interaction to a RAS/RAF/RTK pathway member in InWeb). We compared the degree of copy number gains, and activating SSNVs / indels, in our set of nine genes to 100 matched sets of nine RAS-affiliated genes, showing that the set of nine genes identified through our approach are significantly more enriched for oncogenic copy number gains (*P* = 0.04, using permutation tests, **Supplementary Figure 7**). This analysis confirms that combining NetSig with tumorigenicity experiments is a better approach to identifying driver genes and events in lung adenocarcinomas than naïvely choosing genes in the extended RAS/RAF/RTK pathway.

To allow further exploration of pathway relationships relevant to lung adenocarcinomas the NetSig networks of *AKT2* and *TFDP2* are plotted in **Figs 3f and g**.

## Discussion

Overall our integrated computational and experimental analyses firmly establish network-based approaches can contribute to expanding gene discovery from existing cancer genomes. Not only do the genes in the NetSig5000 set point to new genes in well-established oncogenic pathways (e.g., *AKT2*, *PIK3CB*, *PIK3CG*, *RASGRP1*, *RASGRP3*, **Supplementary Note 6** and **Supplementary Figure 8**), but our results also point to potential new cancer pathways (e.g., *TFDP2* and *MYO7A*, **Supplementary Note 7, Supplementary Figures 9** and **10**, **Supplementary Tables 8** and **9**). For details about the differences between NetSig and other network-based methods, see **Supplementary Note 8, Supplementary Figure 11 and Supplementary Table 10**.

An important feature of NetSig is that it is explicitly designed to disregard any mutation information on the gene being tested so that the signal comes from the genes network alone. This ensures that NetSig *P* values are fully independent of those from existing gene-based statistical test such as MutSig, Oncodrive, GISTIC and RAE (in fact, the MutSig and NetSig *P* values of the same genes are only modestly correlated, Pearson correlation coefficient = 0.05, data not shown). This design choice means that NetSig can be seamlessly combined with (and thus complement) gene-based statistical tests in any computational cancer genome analysis workflow (**Supplementary Note 9** and **Supplementary Figure 12**). The NetSig code is available from www.lagelab.org/resources and the algorithm has been implemented in the FireCloud cancer genome analysis platform (https://software.broadinstitute.org/firecloud/)

NetSig is flexible and can work with many different types of functional genomics network data (**Supplementary Note 9** and **Supplementary Figure 13**). Interestingly, the average genomic inflation when NetSig is run on different sets of transcriptional networks^29^ (i.e., based on data that cannot be effected by ‘knowledge contamination’) is 1.14 and 1.11, respectively (**Supplementary Figures 14** and **15**). This is comparable to the lambda in the protein-protein interaction data when the effect of ‘Cosmic classic’ genes are removed from the analysis (1.09), suggesting that our approach to removing ‘knowledge contamination’ is efficient in canceling out that effect. This strongly suggests that the remaining inflation is due to polygenicity of cancers and not due to any bias or confounders of the NetSig statistic or network data.

While we did not observe any evidence for gain-of-function single nucleotide variants (SSNVs) or insertions or deletions (indels) across *TFDP2* and *AKT2*, it is our expectation that with more samples in the future these genes will be enriched for such events. This is consistent with the observation that NetSig5000 set overall is enriched for genes with lower MutSig *P* values in the Lawrence et al, 2014 set (*P* = 0.04 using a nonparametric two-sample Kolmogorov-Smirnov test). Together, our results strongly suggest that many genes in the NetSig5000 set are likely real intermediate or low frequency driver genes that will reach significance in gene-based statistical tests with more tumor genomes in the future. For a discussion of the study’s limitations, an estimate of how well the NetSig statistic predicts real cancer genes, and information on the benefit of including several cell models and genetic backgrounds in the validation workflow, see the extended discussion in **Supplementary Note 10**.

We expect that with more data in the future the approach we describe here will become increasingly powerful for biological discovery in cancers. We make all results and algorithms available from www.lagelab.org/resources as a resource for the community.

## Contributions

HH developed, benchmarked, and implemented the NetSig algorithm with input from ML and supervision from GG and KL. CRC, YS, ES, NI, EK executed the *in vivo* tumorigenesis experiments with input from HH and KL and supervision from JSB. HH developed and implemented the quantitative analytical framework of *in vivo* tumorigenesis data with input from CRC, YS, ES and supervision from JSB and KL. JDC reanalysed lung adenocarcinoma data with input from HH, JSB, GG and KL. All authors analysed data and discussed the results. HH, WCH, JDC, JSB, GG and KL wrote the manuscript with input from all authors. JSB, GG and KL designed and directed the work. KL initiated and led the study

## Methods

### Calculating the network mutation burden

For a given index gene, NetSig statistic is formalized into a probabilistic score that reflects the index-gene-specific composite mutation burden [i.e. the aggregate of single-gene MutSig suite Q values from^11^] across its first order biological network and is calculated via a three-step process: First, we identify all genes it interacts with directly at the level of proteins, only including high-confidence quality-controlled data from the functional human network InWeb^30,31^ (where the vast majority of connections stem from direct physical interaction experiments at the level of proteins). Second, the composite mutation burden across members of the resulting network is quantified by aggregating single-gene MutSig suite Q values from^11^ into one value ϕ using an approach inspired by Fisher’s method for combining p-values:

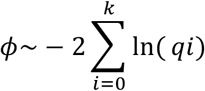

Where p_i_ is the MutSig suite Q value for gene i, and k is the amount of genes in the first order network of the index gene (i.e. the index gene’s degree). Third, by permuting the InWeb network using a node permutation scheme, we compare the aggregated burden of mutations ϕ to a random expectation. In this step, the degree of the index gene, as well as the degrees of all genes in the index gene’s network is taken into careful consideration. The final NetSig score of an index gene is therefore an empirical *P* value that reflects the probability of observing a particular composite mutation burden across its first order physical interaction partners (at the level of proteins) normalized for the degree of the index gene as well as the degrees of all of its first order interaction partners. Because we are interested in estimating the mutation burden independent of the index gene (so that the NetSig results are fully independent of gene-based statistical tests such as MutSig, Oncodrive, GISTIC and RAE), this gene is not included in the analysis and it does not affect the NetSig calculation. This also means that for any given gene MutSig suite significances are independent of NetSig significances (i.e., the Cancer5000 gene set and the NetSig5000 gene set are independently predicted).

### Classifying cancer genes

For each gene represented in InWeb (12,507 or 67% of the estimated genes in the genome), we used the gene-specific NetSig probability to classify it as a cancer candidate gene or not. True positive genes were a set of ‘Cosmic classic’ genes and a set of ‘recently emerging cancer genes’. Specifically, the Cosmic classic set consists of 38 established (or classic) cancer genes from the Catalogue of Somatic Mutations in Cancer (Cosmic, http://cancer.sanger.ac.uk/cosmic, e.g., *TP53*, *BRCA1*, and *BRAF*, **Supplementary Table 1**). The ‘recently emerge cancer genes’ contains 61 genes that have been recently identified as cancer genes from the Sanger Gene Census dataset (http://cancer.sanger.ac.uk/census/, e.g., *MLL2*, *CDK12*, and *GATA2*, **Supplementary Table 1**). The gene set for the purposes of the benchmarking analysis is a set of 87 random genes (**Supplementary Table 1**). True negatives were defined as all genes in InWeb that were not in these three sets which is likely conservative as many of these might be yet undetected cancer genes. We used the NetSig probability as the classifier and calculated the AUC for each gene set. For estimating AUC significances, we generated AUCs for 100 random networks (from Rossin et al., 2011) and calculated the empirical P value.

### Using NetSig to classifying driver genes across 21 tumor types

For each tumor type we calculated tumor-type-specific NetSig scores and classified the corresponding tumor-specific driver genes. For example, we assembled a set of driver genes from breast tumors (BRCA) by identifying genes significantly mutated in this tumor type in^11^. We used mutation data from this tumor type to derive NetSig_BRCA_ scores and measured their classification performance on the BRCA driver genes, which they could accurately distinguish with an AUC = 0.76. We compared this result to the results using NetSig scores derived using the pan-cancer dataset. The pan-cancer NetSig score increased the ability to accurately classify BRCA driver genes slightly to an AUC of 0.77 (for more information see **Supplementary Note 1**, **Supplementary Figure 2**).

### Testing the robustness of the NetSig approach

To test the robustness of the NetSig approach, we tried several alternative permutation methods and calculated the composite mutation burdens of gene networks using both Q and *P* values from the Lawrence et al. 2014 paper. Specifically, to generate the null distribution of network mutation burdens used to assess the significance of observations in the actual data, we both used a node permutation scheme and a full network permutation scheme. Where the node permutation scheme permutes nodes that have similar degree has the advantage of being much faster than the network permutation scheme [explained in detail in^32^], the architecture of the original network is more precisely mirrored in the random networks using the latter method. We ran the full analysis using both approaches and compared the quantile-quantile plots (not shown), and classification of Tiers 1-5 genes. This analysis confirmed that the choice of permutation scheme does not have a major influence on the overall results (**Supplementary Figure 1**). In addition to using q values from^11^ for step 2 in the NetSig calculation (above), we also tried using unadjusted *P* values. For this latter approach the quantile-quantile plots (not shown) as well as the classification of ‘Classic’ and ‘Recently emerging’ cancer genes similar to the results we report in the main text (**Supplementary Figure 1**).

### Generating the NetSig5000 set

We used a node permutation scheme to create 10^6^ permuted networks. NetSig probabilities were determined for every gene in InWeb that was covered by interaction data. The FDR Q values were calculated as described by Benjamini and Hochberg^33^ based on the nominal *P* values controlled for 12,507 hypotheses. We performed NetSig analyses with the pan-cancer Q values, as well as Q values from each of the 21 tumor types for which they were available. As it is a technical limitation of the NetSig approach that it is currently not possible to make 5.5 × 10^6^ network permutations we could not create a dataset where we correct for all 12,500 × 22 hypotheses tested in the NetSig5000 set. For that reason our work does not have the equivalent of the Cancer5000-S (the stringent) set from Lawrence et al.^11^, where the authors control for all hypotheses is carried out simultaneously.

### A multiplexed *in vivo* tumor formation screen in mice

We used the SALE-Y cell model previously described in (Berger et al., 2016) and the HA1E-M cell model previously described in (Kim et al., 2016). Specifically, our earlier work revealed that immortalized small-airway epithelial cells harboring an activating *YAP1* variant are rendered tumorigenic via activation of the EGFR/MAPK pathways [SALE-Y cells^21^] and immortalized kidney epithelial cells harboring an activating *MAPK1* variant are rendered tumorigenic via activation of the PI3K/YAP/NFKB pathways [HA1E-M cells^21–24^]. Briefly, we inserted each gene into barcoded cDNA clones and these clones were transduced into SALE-Y and HA1E-M cells in 96 well plates in arrayed format. The cells were selected with puromycin, expanded, and pooled. Two million cells per pool per site were injected subcutaneously into immunocompromised mice in three sites (interscapular area and left and right flanks) per mouse and tumor formation monitored. The experimental endpoint was reached when any tumor length exceeded 1 cm. Tumors were homogenized and genomic DNA was extracted and sequenced to determine the relative proportion of each inserted DNA barcode. The relative proportions of each barcode serves as a proxy indicating the gene driving a tumor. To deal with data from deploying multiple cell models in parallel on a large set of positive controls, random controls and NetSig candidates we developed a new quantitative analytical framework (below).

### A quantitative analytical framework to compare the tumorigenic potential of NetSig5000 genes to known oncogenes and random genes

We measured the reproducibility and magnitude of the oncogenic signal of the individual gene sets by developing and calculating two complementary metrics: maximum *in vivo* proliferation rate and significance of relative growth:

#### Calculating max proliferation rates

To determine a metric for growth doubling time of cells injected with NetSig5000 genes in the *in vivo* tumors, we calculated the proportion of reads in a tumor normalized to tumor volume and compared to the proportion of reads in the pre injection cell pool where volume for all pooled cells was set to 1 cubic mm (which roughly corresponds to 2 million cells). This was done for all tumors and for each tumor we divided the growth rate with the day the tumor was harvested to normalize for tumor age. This leads to an estimate of the doubling time of the *in vivo* tumor growth for cells driven by overexpression of a particular NetSig candidate. We call this metric max proliferation rate per gene, which is plotted on the x-axis of **Fig. 2b**.

#### Calculating the significance of relative growth

To calculate the significance of relative growth of cells in each cell type (SALE-Y and HA1E-M, respectively) transduced with a particular cDNA clone we plotted the distribution of relative reads in the tumors and compared to the pre injection value. Significances were calculated using a one sided t-test and reported as false discovery rates. We call this metric significance of proliferation rate and plotted the maximum significance (after iterative removal of dominant effects - see below) on the y-axis of **Fig. 2b**.

#### Computational detection of dominant and subjugated oncogenic clones in tumors

When many oncogenic clones are pooled and injected into mice, a single clone often outcompetes other oncogenic clones to dominate the tumor through a highly stochastic process. We refer to outcompeted, but real, oncogenic clones in the tumors as ‘subjugated oncogenic clones’. It is possible to detect subjugated oncogenic clones by iteratively removing dominant clones from the cell pools and repeating the experiments. However, this is very labor intensive. We developed a computational approach where we iteratively removed genes that accounted for more than 50% of the reads in a tumor and repeated the significance of relative growth analysis described above. In **Fig. 2b** we report the best FDR after zero, one or two iterations. We confirmed that the subjugated oncogenic clones detected computationally were indeed driver clones by comparing the results from our computational approach to results from the iterative experimental removal of dominant clones and repetition of the injection of experimentally reduced cell pools into mice from the Berger et al., paper^21^. This analysis showed that genes determined to be significant through our computational iterations also became dominant clones when other dominant clones were first removed from the experimental assay.

### Calculating the sensitivity and specificity of the experimental tumorigenesis assay

#### Sensitivity

We determined how many of the 25 positive control genes were correctly classified as tumor inducing at z-scores of one and two, in both the HA1E-M and SALE-Y model. In the HA1E-M model, 6 genes were classified as tumor inducing at a z score of one and 2 at a z score of two (see **Supplementary Table 7** for details). As we tested a total of 25 genes this gives a sensitivity of 6/25 = 0.24 and 2/25 = 0.08, respectively (**Fig. 2b**). In the SALE-Y model, 7 genes were classified as tumor inducing at a z score of one and 6 at a z score of two giving a sensitivity of 7/25 = 0.28 and 6/25 = 0.24, respectively (**Fig. 2b**). When combining the two assays together the sensitivity increases to 9/25 = 0.36, which is likely because we are testing the tumorigenic potential of genes across several genetic backgrounds. Analogous calculations can be seen for constructs in **Supplementary Table 7**.

#### Specificity

We determined how many of the random genes constructs were correctly classified as non-tumor-inducing at z-scores of one or two (see above), in both the HA1E-M and SALE-Y model. In the HA1E-M model, three genes (*STRADA*, *ZNF346*, and *DRD4*) were classified as tumor inducing at a z score of 1, and one gene (*STRADA*) was classified as tumor inducing at a z score of two. As we tested a total of 79 genes this gives a specificity of 76/79 = 0.96, and 78/79 = 0.99, respectively (**Fig. 2b**). In the HA1E-M model, one gene (*NTRK1*) was classified as tumor inducing at a z score of 1 and z score of 2, respectively. As we tested a total of 79 genes this gives a specificity of 78/79 = 0.99% at both thresholds (**Fig. 2b**). Analogous calculations can be seen for constructs in **Supplementary Table 7**.

### Choosing 25 genes for the validation experiment

We selected the genes based on a number of biological (not being known cancer genes) and technical (available high quality reagents) criteria: <u>First</u>, we selected a set of genes that were either in group 3,4 or 5 of our literature curation groups (meaning they have not already been shown to be cancer genes in humans). <u>Second</u>, we chose the subset of genes for which there were already reagents (meaning open reading frame [ORF] constructs) available from the Genetic Perturbation Platform at the Broad Institute. <u>Third</u>, we chose the set of genes where the ORF constructs had been sequenced and i) did not have any mutations [i.e., that the sequence of the cDNA corresponded to the wild type] and ii) where the sequence of the ORF passed a high quality sequence cutoff to avoid testing ORFs where the sequence of the clone was ambiguous and could have unknown mutations. <u>Fourth</u>, the cell models are optimized for perturbations in certain pathways (i.e., the SALE-Y cells are rendered tumorigenic via activation of the EGFR/MAPK pathways and the HA1E-M cells are rendered tumorigenic via activation of the PI3K/YAP/NFKB pathways). We hypothesized that choosing a set of genes that linked to the pathways activated in each cell model would likely increase the chance to induce tumors in these models. We tested this hypothesis by choosing the 25 genes so, when possible, they interacted directly with members of the pathway activated in the HA1E-M model, but not in the SALE-Y model. However, we see similar validation rates in the two models so it does not seem to have an effect that we are ‘fitting’ the candidates specifically to the HA1E-M model (**Supplementary Note 10**). It is likely that the higher validation rates observed for Netsig candidates (138% of the theoretical expectation, **Discussion**) when using both cell models in parallel is due to a combination of these selection criteria (available reagents and connection to known cancer pathways in the HA1E-M model) and underestimates of the sensitivity of the assay because there is an upper limit to how many true positive oncogenes in a cell pool can induce tumors based on the issues with subjugated clones mentioned above).

### Analysis of oncogene negative lung adenocarcinoma patients

Segmentation was performed using the Circular Binary Segmentation algorithm followed by Ziggurat Deconstruction to infer the length and amplitude of each segment. Recurrent peaks for focal somatic copy number alteration were identified using GISTIC 2.0^8^. A peak was considered to be focally amplified or deleted within a tumor if the GISTIC 2.0–estimated focal copy number ratio was greater than 0.1 or less than −0.1, respectively. Purity and ploidy were estimated using ABSOLUTE^34^. Two peaks were considered the same across tumor types if (i) the known target gene of each peak was the same or (ii) the genomic location of the peaks overlapped after adding 1 mega base to the start and end locations of each gene. For the second criterion, only peaks that contained fewer than 25 genes and were smaller than 10 Mb were considered [for more details see Campbell et al., 2016)]. Because we are executing a case-control analysis of the copy numbers of genes that induce tumors in the SALE-Y model relevant for lung adenocarcinoma our analysis normalizes out any potential effects of, for example gene size, amount of protein-protein interactions a gene has and so forth).

### Data availability

NetSig code, results, and visualizations are available from www.lagelab.org/resources. The protein network data (InWeb version 3.0) is available from www.lagelab.org/resources. Tumor genome data is publicly available from the Lawrence et al, 2014 paper^11^ Lung cancer datasets are available from Campbell et al, 2016^25^. Further data that supports the findings of this study is available from the corresponding author upon request. NetSig is implemented in FireCloud (https://software.broadinstitute.org/firecloud/).

